# Stable Inheritance of Transgene and Yellow Fluorescent Protein Gene Expression in Progeny of Transgenic Cacao (*Theobroma cacao*) Plants

**DOI:** 10.64898/2025.12.09.692773

**Authors:** George Austin, Jesse Jones, Abigail Stevens, Elaine Zhang, Taylor Thompson, Michael Gomez, Geoffrey Vrla, Youngbin Oh, Jean-Philippe Marelli, Carl Jones, Brian Staskawicz, Myeong-Je Cho

**Author notes:** Correspondence: Correspondence (M.-J.C.). (G.A.); (J.J.); (A.S.); (E.Z.); (T.T.); (M.G.); (G.V.); (Y.O.); (B.S.). (J.-P.M); (C.J.).

## Abstract

Genetic engineering tools have the potential to rapidly and precisely improve the genome of slow-to-breed cacao. We previously developed an efficient protocol for transforming cacao using cotyledonary explants derived from secondary somatic embryos via *Agrobacterium tumefaciens*. In this study, we demonstrate that our transformation protocol is successful in elite cultivars, INIAPG-038 and Matina 1-6, producing fertile seeds with stable visual marker inheritance regardless of whether the transgenic plants were used as the pollen or ovule donor. Three vectors were used in the transformations, each containing genes for *enhanced yellow fluorescent protein* (*eyfp*) and *neomycin phosphotransferase* II (*npt*II). Three transgenic INIAPG-038 events and one transgenic Matina 1-6 event were used to evaluate seed fertility and the stability of transgene inheritance in cacao seeds and plants. The T1 progeny of these four transgenic events were analyzed for YFP expression and transgene presence. YFP expression segregated at a 1:1 ratio in all events when the transgenic plants were crossed with non-transgenic plants, while a 3:1 segregation was observed when transgenic events were crossed with each other. The transgenic plants exhibited a normal phenotype compared to non-transgenic control plants, producing seeds with a 97% germination rate.

## 1. Introduction

The global multi-billion-dollar chocolate industry is based on the fruit of cacao (*Theobroma cacao L.*) trees. Cacao is predominantly cultivated in West Africa, followed by Southeast Asia and South America, where it was originally domesticated [1]. There are ten recognized genetic groups of cacao [2]. The most flavorful groups, Criollo and Nacional, suffer from susceptibility to disease and pests [3], as well as an accumulation of yield-impacting deleterious mutations [4]. Despite a price premium for Criollo varieties with floral, fruity notes, highly productive Amelonado hybrids dominate the global market. Globally, cacao production is greatly reduced by diseases, whether affecting Criollo and Nacional in South America or Amelonado in West Africa [5]. Developing new cacao cultivars that combine high yield, quality, and resistance to diseases and pests among different genetic groups using traditional breeding methods is a time-consuming and complex process.

As is the case with most fruit tree species, traditional plant breeding of cacao can take 20 or more years to release a new cultivar [6] and is hampered by lengthy juvenile periods, high heterozygosity, self-incompatibility, and a lack of funding [7,8]. On the other hand, genetic modification using *Agrobacterium* and biolistic methods, which allows for the addition of valuable genes not naturally present within a species, has achieved dramatic crop improvements such as resistance to disease, pests, abiotic stress, and herbicide, along with enhanced nutrient use efficiency, and improved photosynthesis, ultimately resulting in increased yields [9]. This approach overcomes the challenges associated with traditional breeding. In addition, recent genome editing technology enables precise modification of genes or regulatory sequences already present within a species, providing an expanded toolkit for trait development as well as genetic discovery. These genome modifications can alter expression levels, facilitate the discovery of novel alleles, manipulate quantitative trait loci [10], and generate knockout lines for studying gene function. The first stable transformation of cacao occurred in the PSU SCA-6 cultivar, resulting in the production of six pods containing 143 transgene-positive and 139 transgene-negative seeds from a single transgenic event [11]. In a previous study [12] we reported a successful transformation of INIAPG-038, a high-yielding elite breeding line developed by Mars Wrigley, Incorporated, in collaboration with the United States Department of Agriculture (USDA) and Instituto Nacional de Investigaciones Agropecuarias (INIAP). This line also has some resistance to fungal diseases *Moniliophthora roreri* (frosty pod) [13], and *Moniliophthora perniciosa* (witches’ broom) [14] which are widespread in Central and South America [15]. Matina 1-6, which has been sequenced, belongs to the most widely planted Amelonado genetic group [16]. It is susceptible to cacao swollen shoot virus (CSSV) [17] which is prevalent in West Africa where it is commonly cultivated. There are no previous reports in the literature of successful transformation in Matina 1-6.

Although genetic transformation can be a powerful tool, unintended detrimental changes may take place during the tissue culture and transformation process. Mutations, including those causing sterility, can arise due to somaclonal variation, where stress during tissue culture can cause DNA methylation, activation of transposable elements, or the creation of new mutations [18,19]. The expression of the inserted genes may be affected by nearby endogenous genes, complex transgene loci, and excessive expression [20]. Inserted genes may interfere with important endogenous genes that are located nearby, bisected by, or homologous to the transgene [21–23]. Transgene silencing can also be an issue, occurring at the transcriptional level through DNA methylation or post-transcriptionally through RNA degradation [24]. Due to these potential issues as well as chimerism and multiple copy insertions, the transgene inheritance ratio expected when the T0 is outcrossed may skew away from the expected Mendelian segregation ratio. It is crucial to screen novel transgenic lines for all these issues through T0 and T1 plant analysis to ensure the genetic transformation was successful. In cacao, germplasm is commonly disseminated through polyclonal seed gardens, producing hybrid seed in West Africa and through clonal cuttings in the Americas and Asia [25]. In both instances, stable expression and inheritance of transgenes are important for creating new genetically engineered cultivars or breeding lines. Once stably integrated, transgene inheritance and the stability of mRNA and protein expression levels will remain constant over generations, even when backcrossed or outcrossed [26].

In this study, we demonstrate that our genetic transformation protocol for 2 cacao cultivars, INIAPG-038 and Matina 1-6, is successful in producing plants with stably incorporated transgenes and stable transgene expression. Importantly, the transformation process did not adversely affect seed fertility, pollen and ovule viability, or plant phenotype. These findings were confirmed by evaluation of plant morphology, cross-pollination compatibility, seed size, germination rate, and transgene expression and inheritance in both T_0_ and T_1_ plants.

## 2. Results and Discussion

The primary goal of this research is to confirm whether transgenic cacao plants generated using our protocol [12] are healthy, phenotypically normal, fertile, and capable of transmitting inserted transgenes in a Mendelian fashion without any adverse effects. To evaluate stable transgene expression and inheritance, 3 independent transgenic events of INIAPG-038 and 1 transgenic event of Matina 1-6 were analyzed. One event (INIAPG-038 EVT1) was transformed with the pDDNPTYFP-1 vector (Figure 1A), while 2 events (INIAPG-038 EVT2 and EVT3) were transformed with the pDDNPTYFP-2 vector (Figure 1B), all 3 were generated previously via *Agro-bacterium*-mediated transformation [12]. In contrast, Matina 1-6 EVT1 was generated for this study using the pMGCC3YFP vector, which contains *cas9* and gRNA cassettes in addition to *npt*II and *eyfp* gene cassettes (Figure 1C), following the same transformation protocol [12].

**Figure 1.**
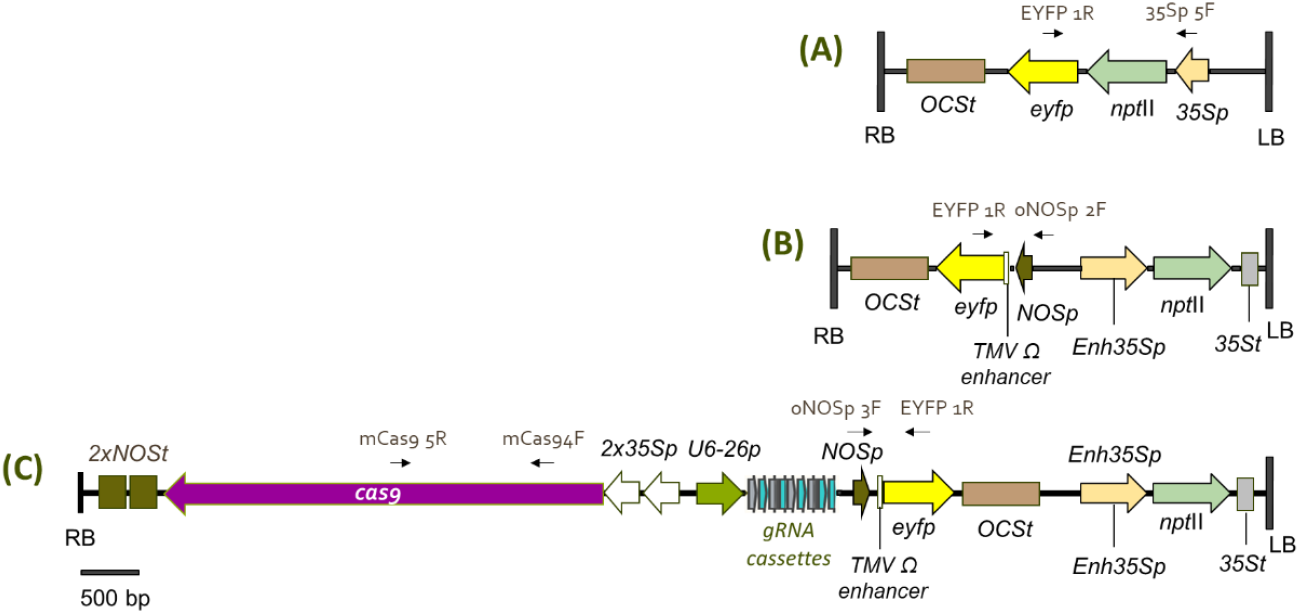
Schematic diagram of three transformation vectors used for *Theobroma cacao* L. transformation. (A) pDDNPTYFP-1 contains a 3,637-bp T-DNA fragment with a translational fusion of the neomycin phosphotransferase II (*npt*II) and enhanced yellow fluorescent protein (*eyfp*) gene, driven by a CaMV 35S promoter. (B) pDDNPTYFP-2 contains a 4,416-bp T-DNA fragment with two gene cassettes: *eyfp* is driven by a Nos promoter/TMV Ω enhancer, and *npt*II is driven by an enhanced CaMV 35S promoter. (C) pMGCC3YFP contains a 11,460-bp T-DNA fragment with four gene cassettes: *eyfp* is driven by a Nos promoter/TMV Ω enhancer, *npt*II is driven by an enhanced CaMV 35S promoter, and two additional cassettes encode *cas9* and *gRNA*.

Transformation frequencies differed markedly between the 2 cultivars: INIAPG-038 produced 3 independent events from 82 secondary somatic embryos (3.7% at the T_0_ plant level) [12], whereas Matina 1-6 yielded 1 event from 162 embryos (0.62%). The lower transformation frequency observed in Matina 1-6 may be attributable to its reduced efficiency in secondary somatic embryogenesis and T-DNA delivery [12], as well as differences associated with the larger T-DNA region (Figure 1). These results underscore the genotype-dependent variability in cacao transformation and the impact of vector composition on transformation frequency.

Unique primer sets were designed for each construct (Table S1) and validated through polymerase chain reaction (PCR) analysis using plasmid DNA and genomic DNA extracted from leaf tissues of non-transgenic plants and transgenic events (Figure 2). The presence of the *eyfp* transgene was confirmed by PCR using primer sets, 35Sp 5F/EYFP 1R, oNOSp 2F/ EYFP 1R, and oNOSp 3F/EYFP 1R for pDDNPTYFP-1, pDDNPTYFP-2 and pMGCC3YFP vectors, respectively (Figure 1), as well as their corresponding transgenic events, INIAPG-038 EVT1 (Figure 2A), INIAPG-038 EVT2/EVT3 (Figure 2B) and Matina 1-6 EVT1 (Figure 2C). For Matina 1-6 EVT1, the *cas9* transgene was also confirmed using the mCas9 4F/mCas9 5R primer set (Figure 2D). A 1,281-bp amplified product corresponding to the CaMV 35S promoter::*npt*II::*eyfp* gene fragment was detected in the genomic DNA of INIAPG-038 EVT1 using the 35Sp 5F/ EYFP 1R primer set (Figure 2A; Table S1), while a 593-bp NOS promoter::TMV Ω enhancer::*eyfp* gene fragment was amplified from INIAPG-038 EVT2 and INIAPG-038 EVT3 using the oNOSp 2F/EYFP 1R primer set (Figure 2B; Table S1). In Matina 1-6 EVT1, a 594-bp NOS promoter::TMV Ω enhancer::*eyfp* gene fragment was amplified using the oNOSp 3F/EYFP 1R primer set (Figure 2C; Table S1), and/or a 1,235-bp *cas*9 fragment was amplified using the mCas9 4F/mCas9 5R primer set (Figure 2D; Table S1).

**Figure 2.**
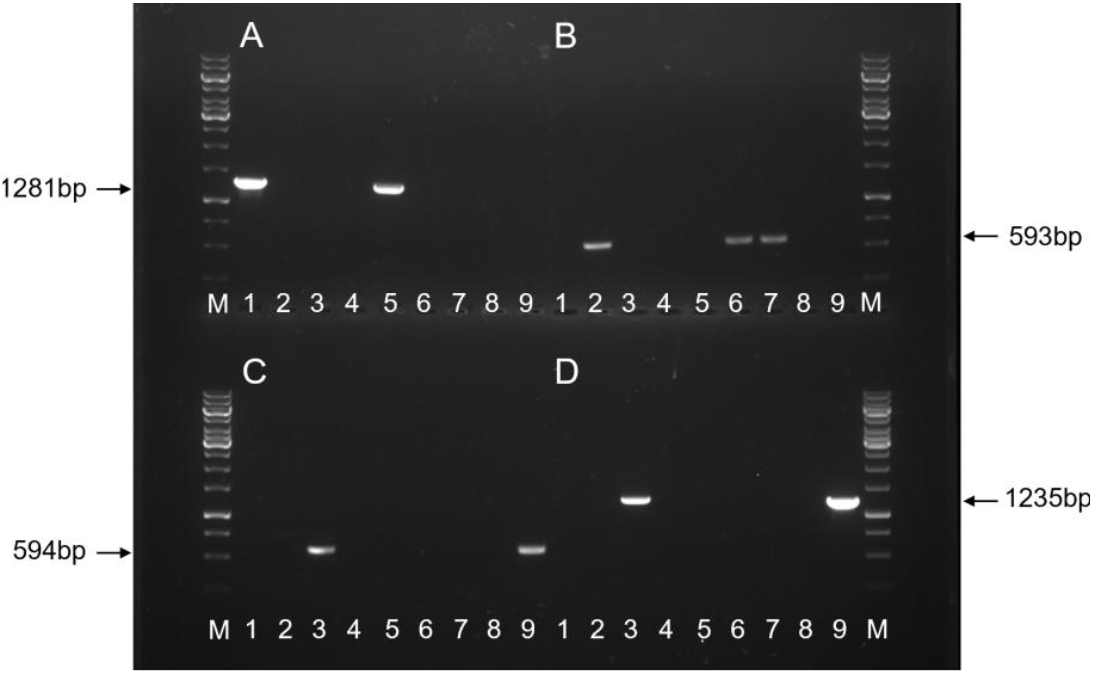
PCR analysis of genomic DNA from four different transgenic *Theobroma cacao* L. events. Genomic DNA was extracted from leaf tissues of one non-transgenic INIAPG-038 control, three transgenic INIAPG-038 events (EVT1, EVT2 and EVT3), one non-transgenic Matina 1-6 control, and one transgenic Matina 1-6 event (EVT1). M, GeneRuler 1-kb DNA ladder; Lane 1, pDDNPTYFP-1 plasmid; Lane 2, pDDNPTYFP-2 plasmid; Lane 3, pMGCC3YFP plasmid; Lane 4, non-transgenic INIAPG-038; Lane 5, INIAPG-038 EVT1; Lane 6, INIAPG-038 EVT2; Lane 7, INIAPG-038 EVT3; Lane 8, non-transgenic Matina 1-6; Lane 9, Matina 1-6 EVT1. (A) Amplified 1,281-bp fragment products using 35Sp 5F and EYFP 1R. (B) Amplified 593-bp fragment products using oNOSp 2F and EYFP 1R. (C) Amplified 594-bp fragment products using oNOSp 3F and EYFP 1R. (D) Amplified 1,235-bp fragment products using mCas9 4F and mCas9 5R.

All four T_0_ events exhibited YFP expression in both leaf and root tissues (Figure S1). However, visual YFP expression driven by the CaMV35S promoter in INIAPG-038 EVT1 was notably stronger than that of the other 3 events—INIAPG-038 EVT2, EVT3, and Matina 1-6 EVT1—where expression was driven by the NOS promoter with the TMV Ω enhancer (Figure S1). Western blot analysis showed that YFP protein levels driven by the CaMV35S promoter were also 10-to 30-fold higher than those driven by the NOS promoter with the TMV Ω enhancer (Figure S2). This is similar with findings by Sanders et al. [27], who reported that the CaMV35S promoter facilitated a 30-fold higher expression of the *npt*II gene in petunias compared to the NOS promoter. Although the TMV Ω enhancer has also been shown to increase transgene expression in plants by 3% when attached to CaMV35S promotor [28] and by 100% when attached to heat shock protein (*hsp80*) promoter [29], in our study, the NOS promoter with the TMV Ω enhancer did not surpass the YFP expression levels of the CaMV35S promoter (Figure S1). In INIAPG-038 EVT2 and EVT3, YFP expression highlights the leaf veins, making them clearly visible, whereas in Matina 1-6 EVT1, the YFP expression is more evenly distributed between veins and interveinal areas (Figure S1). Since all three events share the same NOS promoter::TMV Ω enhancer, the observed differences in YFP expression are likely attributable to variations in integration site, copy number, or structural modifications to the promoter or terminator regions.

To evaluate seed fertility and transgene expression and inheritance in progeny plants, transgenic cacao flowers were hand-pollinated with transgenic or non-transgenic counterparts using plants approximately 3 to 4 years post-transformation. T_0_ plants were phenotypically normal compared to non-transgenic plants (Figure 3A). Stamens were removed within twenty-four hours of flower opening using fine point tweezers, and the anthers were rubbed directly onto the stigma of the ovule donor, with at least two anthers used per flower. Successful pollination was indicated by a lack of flower senescence 48 hours post-opening. Flowers from incompatible crosses senesced within 5 days post-pollination (Figure 3B), while compatible crosses resulted in pods ripening approximately 180 days post-pollination (Figures. 3C, D and E). In nature, insects play a key role in cacao pollination, particularly for self-incompatible plants [30]. Self-pollination of INIAPG-038 did not produce any pods due to self-incompatibility, while self-pollination of Matina 1-6 yielded pods, regardless of whether the plants were transgenic or non-transgenic (Table 1). Self-incompatibility, common in many cacao cultivars, involves 2 separate processes of gamete selection [31]. Matina 1-6 is from the Amelonado genetic group, which is known for having alleles conferring self-compatibility [16], while INIAPG-038 has a mixed heritage, including the Nacional group which is known to have a low-incidence of self-compatibility alleles [31]. In this study, self-compatibility in non-transgenic Matina 1-6 and self-incompatibility in non-transgenic INIAPG-038 were unaffected by transformation (Table 1).

**Table 1.**
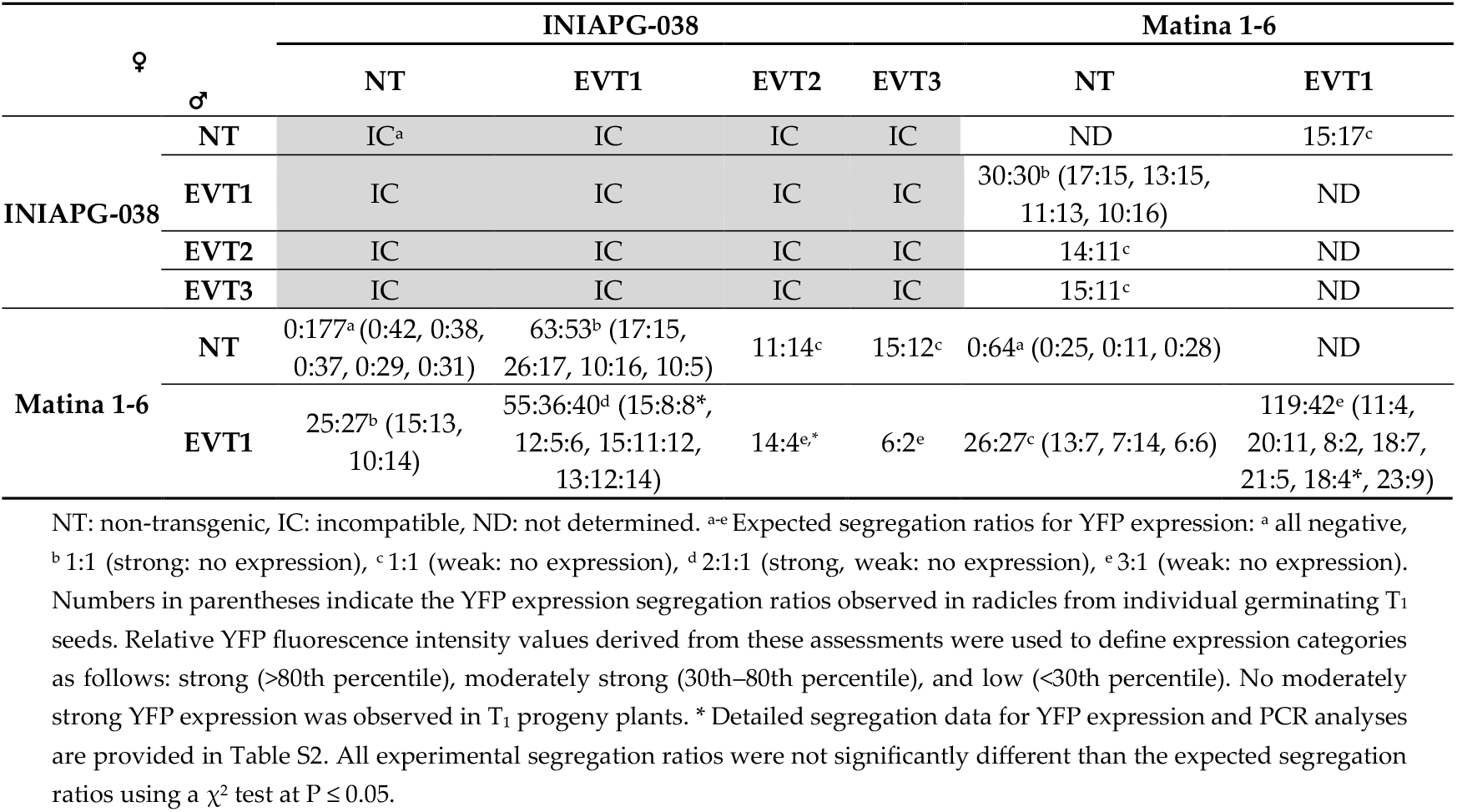
Yellow fluorescence protein (YFP) expression segregation from inter- and intra-crosses among transgenic and non-transgenic cacao plants.

**Figure 3.**
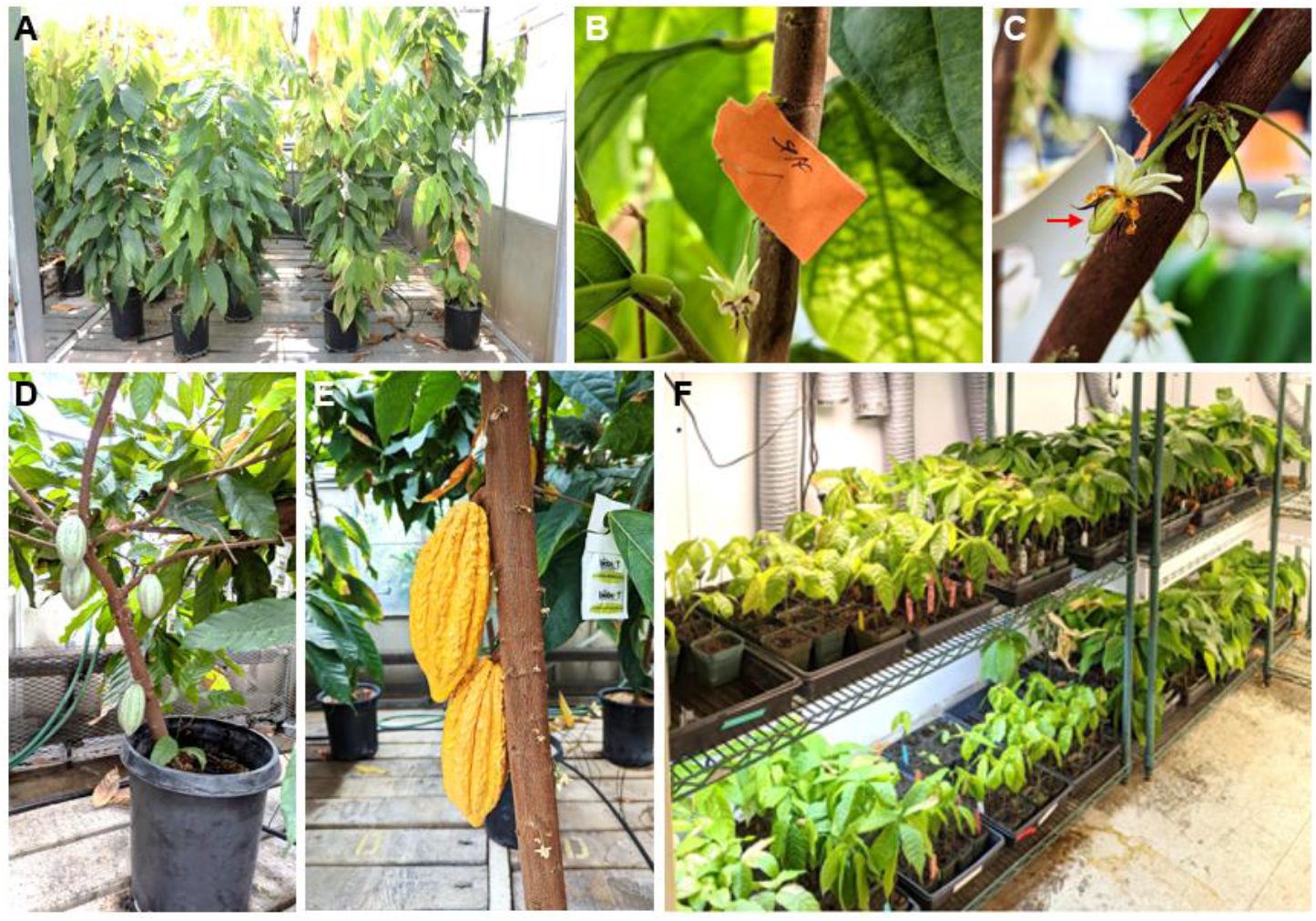
T_0_ and T_1_ cacao plants, flowers, fruits, and germination test used in this study. (A) Non-transgenic plants (left two rows) and T_0_ INIAPG-038 plants (right two rows). (B) Flowers three days after self-pollination on a non-transgenic INIAPG-038. (C) A fruit six days after pollination on a non-transgenic Matina 1-6 ovule pollinated with non-transgenic INIAPG-038 pollen. (D) Fruits from self-pollinated Matina 1-6 EVT1 approximately three months after pollination. (E) Fruits resulting from a cross between INIAPG-038 EVT1 and Matina 1-6 EVT1. (F) Germination test using T_1_ seeds from crossed fruits.

Differences in the expression of YFP among T_1_ seeds and plants were observed across the transformation vectors, as expected. Transgenic progeny of INIAPG-038 EVT1, which utilized the CaMV 35S promotor for YFP, exhibited a very strong YFP expression in all tissue types. In the leaf and root tissues of T_1_ plants from crosses of non-transgenic INIAPG-038, INIAPG-038 EVT1, INIAPG-038 EVT2, INIAPG-038 EVT3, and Matina 1-6 EVT1 with non-transgenic Matina 1-6 (Figure 4A), the level and pattern of YFP expression were the same as those of individual T_0_ counterparts (Figure S1). INIAPG-038 EVT1 transgenic progeny, carrying the CaMV 35S promoter for YFP, exhibited significantly stronger YFP expression compared to the T_1_ progeny of INIAPG-038 EVT2, EVT3, and Matina 1-6 EVT1, which utilize the NOS promoter/TMV Ω enhancer for YFP (Figure 4A). In the seed of INIAPG-038 EVT1, YFP expression was visible after manually removing the fruit pulp and seed coat or more rapidly by slicing the seed in half (Figure 4B). In contrast, seeds from crosses with INIAPG-038 EVT2, EVT3, and Matina 1-6 EVT1, all of which had the (NOS) promotor::TMV Ω enhancer for YFP, did not show visible YFP expression in the kernel but exhibited expression in the radical and first true leaves after seed germination. Strong YFP expression was observed in a YFP-positive seed resulting from a cross with a non-transgenic Matina 1-6 ovule donor (Figure 4B). Seeds from a cross between a transgenic INIAPG-038 EVT1 ovule donor and a non-transgenic Matina 1-6 pollen donor (Figure 4C) also exhibit YFP expression, but the YFP expression in the pulp surrounding the seeds is determined by the source of the ovule donor plant (Figures 4B and C).

**Figure 4.**
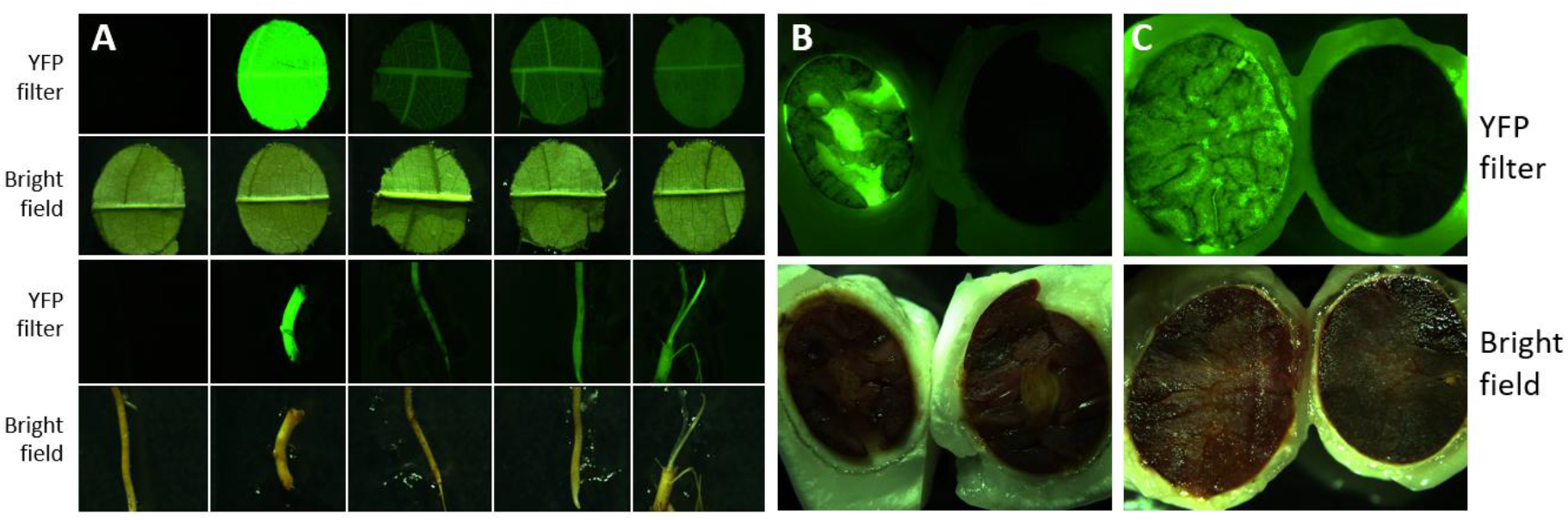
YFP expression in selected tissue of progeny from crosses made between non-transgenic and T_0_ *Theobroma cacao* L. plants. (A) Top row and third row: YFP filter; second row and bottom row: brightfield. From left to right: progeny leaf and root samples from crosses of non-transgenic INIAPG-038, INIAPG-038 EVT 1, INIAPG-038 EVT 2, INIAPG-038 EVT 3, and Matina 1-6 EVT1 with non-transgenic Matina 1-6. (B) Seeds from a cross between a non-transgenic Matina 1-6 ovule donor and a transgenic INIAPG-038 EVT1 pollen donor, showing a transgenic seed (left) and a non-transgenic seed (right). (C) Seeds from a cross between transgenic INIAPG-038 EVT1 ovule donor and a non-transgenic Matina 1-6 pollen donor, showing a transgenic seed (left) and a non-transgenic seed (right). The pulp surrounding the seeds seen in (C) express YFP, consistent with the transgenic ovule donor plant.

YFP expression in T_1_ progeny tissues followed expected Mendelian segregation ratios for all crosses (Table 1). PCR analysis of progeny from selected crosses was consistent with observations from visual inspections (Table S2). Crosses between transgenic and non-transgenic cacao plants from all 4 events demonstrated a 1:1 segregation ratio of YFP expression (Tables 1), consistent with previous findings by Maximova et al. [11]. Copy-number analysis using ddPCR indicated that Matina 1-6 EVT1 and INIAPG-038 EVT1 each contain a single copy of *eyfp*, whereas INIAPG-038 EVT2 and EVT3 contain 3 copies (Figure 5), confirming that all these 4 events in our study have single transgene integration sites. Self-pollination of Matina 1-6 EVT1 yielded a 119:42 segregation, corresponding to the expected 3:1 ratio of YFP-expressing to non-YFP-expressing T_1_ progeny plants (Table 1). Visual YFP expression levels were low and similar between T_1_ progeny fixed (biallelic) for the transgene and those still segregating for it (Tables 1 and S2).

**Figure 5.**
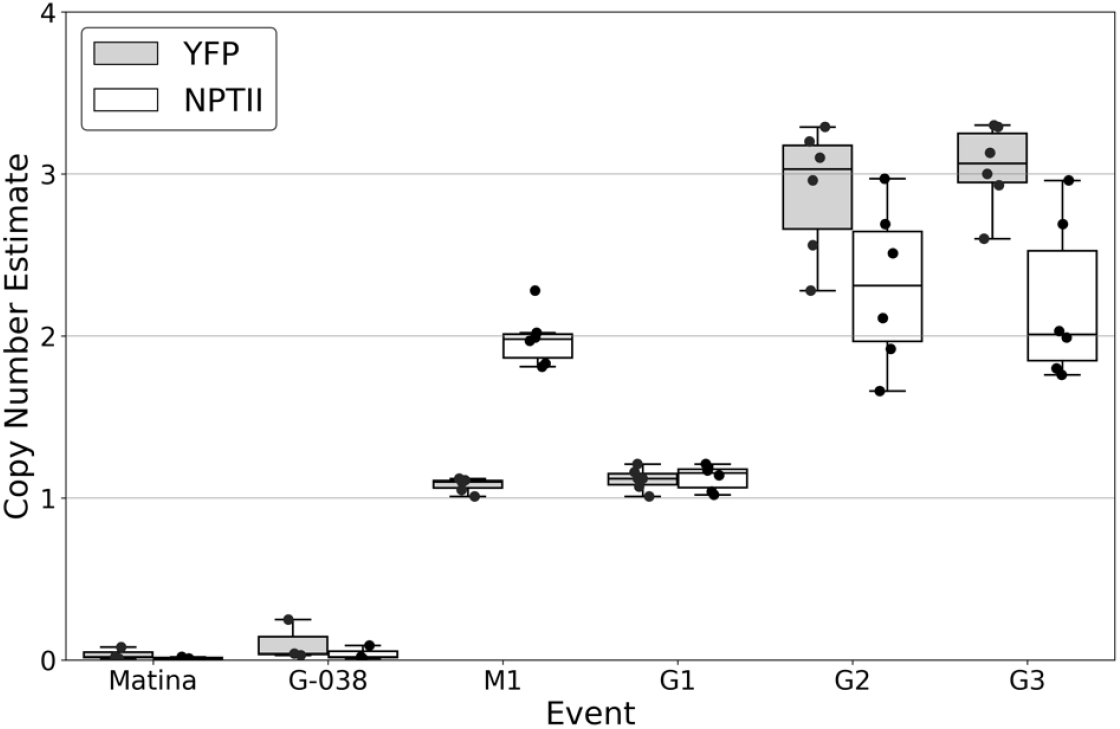
Copy number estimate of YFP and NPTII in T_0_ *Theobroma cacao* L. plants. Calculations of YFP and NPTII copy number performed using the Bio-Rad QX200 digital droplet PCR (ddPCR) system and QuantaSoft Analysis software. Copy number estimate refers to the target counts of the transgene divided by 0.5 x target counts of the TcEF1a reference gene. Values are mean and interquartile range of three separate measurements of two leaf samples. Matina, non-transgenic Matina 1-6; G-038, non-transgenic INIAPG-038; M1, transgenic Matina 1-6 EVT1; G1, transgenic INIAPG-038 EVT1; G2, transgenic INIAPG-038 EVT2; G3, transgenic INIAPG-038 EVT3.

Reciprocal crosses between INIAPG-038 EVT1 and Matina 1-6 EVT1 produced a segregation of 55:36:40 (strong: weak: no expression). Although this pattern did not follow an expected 1:1:1:1 ratio (strong: moderately strong: weak: no expression), no progeny plants with moderately strong expression were observed, and it instead reflected a 2:1:1 (strong: weak: no expression) ratio for expression level because the strong YFP expression driven by the CaMV 35S promoter in INIAPG-038 EVT1 overshadowed the contribution of the weaker NOS promoter::TMV Ω enhancer. As a result, progeny inheriting both transgenes displayed expression levels indistinguishable from those inheriting only the CaMV 35S-driven YFP (Tables 1 and S2). Nevertheless, the overall positive-to-negative YFP segregation still aligned with the expected 3:1 ratio (Tables 1 and S2). Crosses between INIAPG-038 EVT2 or EVT3 and Matina 1-6 EVT1, all expressing YFP under the NOS promoter::TMV Ω enhancer, (demonstrated segregation ratios of 14:4 and 6:2, respectively, both consistent with a 3:1 pattern and characterized by uniformly weak expression (Tables 1 and S2). In these crosses, visual YFP expression was comparable between progeny inheriting a single NOSp::Ω-driven transgene and those inheriting NOSp::Ω-driven transgenes from both parental events (Table S2), reflecting the low YFP expression levels of the parental events (Figure S1).PCR analysis of the INIAPG-038 EVT1 × Matina 1-6 EVT1 and INIAPG-038 EVT2 × Matina 1-6 EVT1 crosses showed a 1:1:1:1 segregation ratio (Table S2). For the INIAPG-038 EVT1 × Matina 1-6 EVT1 cross, 5 plants amplified CaMV 35S promoter::*nptII::eyfp*, NOS promoter/TMV Ω enhancer/*eyfp* (oNOSp 3F/EYFP 1R) and cas9 fragments, 10 amplified only CaMV 35S promoter::*nptII::eyfp*, 8 amplified NOS promoter/TMV Ω enhancer/*eyfp* and *cas9*, and 8 showed no amplication. Because plasmid pMGCC3YFP used to generate Matina 1-6 EVT1 carries the NOSp::Ω::eyfp–cas9 cassette (Figure 1C), plants with only these fragments displayed weak YFP expression (Table S2). Visual determination of these tested plants revealed 15 plants with strong YFP expression driven by the CaMV 35S promoter from INIAPG-038 EVT1 with or without Matina 1-6 EVT1, 8 with weak expression driven by the NOS promoter/TMV Ω enhancer from Matina 1-6 EVT1, and 8 with no expression (Table S2). For the INIAPG-038 EVT2 x Matina 1-6 EVT1 cross, 5 plants showed amplification of both NOS promoter/TMV Ω enhancer/*eyfp* fragments using either (oNOSp 2F/EYFP 1R and oNOSp 3F/EYFP 1R) plus *cas9*; 4 amplified the oNOSp 2F/EYFP 1R fragment only; 5 amplified the oNOSp 3F/EYFP 1R fragment and cas9; and 4 showed no amplification (Table S2). Visual results for these tested plants indicated that 14 YFP-positive plants containing NOS promoter/TMV Ω enhancer/*eyfp* and/or *cas9*, and 4 YFP-negative plants. In self-pollinated Matina 1-6 EVT1, PCR analysis revealed 18 transgene-positive progeny and 4 negative progeny, corresponding to a 3:1 segregation ratio. These results are consistent with Mendelian inheritance of a single transgene locus.

Genetic transformation in plants can lead to unintended effects, including male sterility or low seed set, due to somaclonal variation and disruptions caused by the transformation process. Tissue culture and transformation process may introduce genetic and epigenetic changes that affect fertility [32,33], while transgene insertion or mutagenesis can disrupt endogenous genes critical for reproduction, leading to sterility or developmental abnormalities [34]. Male sterility can also result from disruptions to genes regulating pollen development. Additionally, overexpression of stress-related genes or disruption of hormonal pathways, such as auxins, gibberellins or abscisic acids, can impair reproductive processes and seed development, resulting in low seed set [35,36]. However, in our study, all four transgenic cacao events set fruits and seeds without affecting seed set percentages, regardless of the transgenic nature of the pollen or ovule donor (Table 1). Additionally, T_1_ seed sizes of transgenic and non-transgenic plants were not significantly different (Table 2).

**Table 2.** Germination rates and size of seeds from individual cacao pods produced from non-transgenic plants and crosses between transgenic and non-transgenic plants.

| Ovule donor | Pollen donor | Germination Rate | Average seed length (cm)* | Average seed width (cm)** |
| --- | --- | --- | --- | --- |
| INIAPG-038 | Matina 1-6 | 31/31 | 1.78 | 0.83 |
| INIAPG-038 | Matina 1-6 | 29/31 | 2.08 | 1.03 |
| INIAPG-038 | Matina 1-6 | 37/38 | 2.06 | 1.13 |
| INIAPG-038 | Matina 1-6 EVT 1 | 24/24 | 2.24 | 1.15 |
| INIAPG-038 EVT1 | Matina 1-6 EVT 1 | 38/38 | 2.36 | 1.16 |
| INIAPG-038 EVT1 | Matina 1-6 EVT 1 | 39/39 | 2.10 | 1.10 |
| INIAPG-038 EVT1 | Matina 1-6 EVT 1 | 27/27 | 2.20 | 1.07 |
| INIAPG-038 EVT2 | Matina 1-6 EVT 1 | 22/23 | 2.18 | 1.07 |
| INIAPG-038 EVT3 | Matina 1-6 EVT 1 | 8/12 | 2.41 | 1.15 |
| Matina 1-6 | Matina 1-6 | 11/12 | 2.18 | 1.13 |
| Matina 1-6 | INIAPG-038 EVT1 | 25/26 | 2.13 | 1.05 |
| Matina 1-6 | Matina 1-6 EVT 1 | 21/22 | 1.89 | 1.06 |
| Matina 1-6 | Matina 1-6 EVT 1 | 20/20 | 1.99 | 1.06 |
| Matina 1-6 | Matina 1-6 EVT 1 | 12/12 | 2.07 | 1.02 |
| Matina 1-6 EVT 1 | Matina 1-6 EVT 1 | 31/32 | 1.89 | 0.91 |
| Average |  | 375/387 (97%) | 2.10 | 1.06 |

\* and \*\* indicate no statistical difference (ANOVA, $p > 0.05$ ) by column; PSU used a single event.

Seeds produced by the crosses in this study germinated and formed phenotypically normal plants at a rate of 97% (Figure 3F, Table 3). Most pods harvested in this study contained a couple of seeds noticeably smaller than average; these also formed normal plants but took longer to do so. There were also a negligible number of undeveloped seed structures found within harvested pods that were not included in our results. As the entire cacao industry is based on the cacao seed yield, it is of utmost importance that seeds are not negatively affected as a result of the transformation process. Seed size appears to be unaffected by the transgenic nature of either INIAPG-038 or Matina 1-6 compared to non-transgenic null-segregant seed (Table 3). There also appears to be no correlation between transgene inheritance and germination rates or seed size within pods (Table 3), similar to results in transgenic pear seeds [37]. It is unlikely that seeds inheriting the transgene would be smaller as that would require the genomic imprinting to be affected by the insertion of the transgene or the tissue culture process [38].

**Table 3.** Distribution of seed sizes within single pods of cacao by transgene inheritance based on expression of yellow fluorescence protein (YFP).

| Ovule donor | Pollen donor | Inheritance ratio | YFP from INIAPG-038 EVT1 with or without Matina 1-6 EVT1*** |  | YFP from only Matina 1-6 EVT1 (heterozygous) *** |  | No YFP from either parent (null segregant)*** |  |
| --- | --- | --- | --- | --- | --- | --- | --- | --- |
|  |  | YFP expression segregation ratio* | Length (cm)** | Width (cm)** | Length (cm) | Width (cm) | Length (cm) | Width (cm) |
| INIAPG-038 EVT1 | Matina 1-6 EVT1 | 15:11:12 | 2.42 | 1.19 | 2.41 | 1.19 | 2.26 | 1.09 |
| INIAPG-038 EVT1 | Matina 1-6 EVT1 | 11:8:8 | 2.10 | 1.00 | 2.36 | 1.19 | 2.19 | 1.04 |
| Matina 1-6 | Matina 1-6 EVT1 | 0:7:14 | N/A | N/A | 2.00 | 1.15 a | 1.84 | 1.04 b |
| Matina 1-6 | Matina 1-6 EVT1 | 0:13:7 | N/A | N/A | 2.06 | 1.14 a | 1.86 | 0.96 b |

\* Number of seeds in a pod with high YFP expression, moderate YFP expression, or no YFP expression. \*\*Means followed by the same or no letter indicate no significant difference (ANOVA, $p > 0.05$ ) compared to corresponding measurements in other columns of a row. \*\*\* Visual YFP expression was evaluated in the radicles of individual germinating T<sub>1</sub> progeny seeds.

In conclusion, transgenic cacao plants generated from the elite cultivars INIAPG-038 and Matina 1-6 using our transformation protocol exhibited no detectable adverse phenotypic differences relative to their non-transgenic counterparts. Furthermore, segregation ratios for YFP expression confirmed the stable inheritance of the inserted transgenes. This transformation protocol thus provides a reliable approach for cacao improvement through genetic modification and gene editing techniques

## 4. Materials and Methods

### 4.1. Plant Materials

Transgenic and non-transgenic cacao plants of INIAPG-038 and Matina 1-6 used in this study were generated using tissue culture and genetic transformation, as previously described by Jones et al. [12]. The donor material for these cultivars was shipped from USDA ARS Subtropical Horticulture Research Station in Miami, FL. Three independent transgenic INIAPG-038 events were produced in a previous study; 1 event (INIAPG-038 EVT1) transformed with pDDNPTYFP-1 (Figure 1A), and 2 events (INIAPG-038 EVT2 and INIAPG-038 EVT3) with pDDNPTYFP-2 (Figure 1B). The transgenic Matina 1-6 event (Matina EVT1) was generated using the pMGCC3YFP vector (Figure 1C) according to the protocol published by Jones et al. [12].

In total, for each non-transgenic plant, there were 2-3 trees. For transgenic Matina 1-6 EVT 1 and INIAPG-038 EVT 1, 2, and 3, there were 12, 11, 3, and 3 trees, respectively. At the time of pollination for this study, transgenic INIAPG-038 trees were approximately 4 years post transformation and approximately 3 years since embryo germination, and transgenic Matina 1-6 trees were one year younger. Non-transgenic cacao trees were ∼ 4 years old at the time of pollination.

### 4.2. Transformation Vectors

Three binary vectors, pDDNPTYFP-1, pDDNPTYFP-2 and pMGCC3YFP, each containing *eyfp* and *npt*II, were used for cacao transformation. pDDNPTYFP-1 contains a neomycin phosphotransferase II (*npt*II)-GlyLink-enhanced yellow fluorescent protein (*eyfp*) translational fusion, driven by the cauliflower mosaic virus 35S (CaMV 35S) promoter (35Sp) (Figure 1), [12] while pDDNPTYFP-2 contains 2 separate gene cassettes, *npt*II driven by the enhanced 35Sp and *eyfp* driven by the Nos promoter (NOSp)/TMV Ω enhancer (Figure 1B), [12]. Both vectors were built in the pCAMBIA2300 backbone. For pMGCC3YFP (Figure 1C), the pCAMBIA2300 binary vector backbone was assembled with a CRISPR expression system as described in Gomez et al [39]. Subsequently, the YFP expression cassette, consisting of a NOSp/ NOSp/TMV Ω enhancer, *eyfp*, and OCS terminator, was PCR-amplified using Phusion HF Polymerase [New England Biolabs (NEB), Ipswich, MA] and cloned into the *Kpn*I site of the binary vector by Gibson Assembly (NEB, Ipswich, MA).

### 4.3. Plant Care

Plantlets were transferred from tissue culture into 10-cm pots filled with Sun Gro™ potting mix (Sun Gro Horticulture Distribution Inc., Agawam, MA) when they had developed 8-12 leaves. Plants were maintained in a growth chamber at 28℃, 80% RH, 16-hour photoperiod at 100-250 μmol m^-2^ s^-1^ produced by fluorescent bulbs for 1 month before being transferred to the greenhouse. In the greenhouse, plants were grown under white 40% netting shade cloth and repotted as needed, with final transplantation to 10-gallon pots. Greenhouse conditions were maintained at a minimum temperature of 20℃ without supplemental lighting. The plants were pruned to maintain a single trunk architecture, kept at a height of ∼2.2 m, and spaced at 1.6 m^2^ per plant. Plants were automatically fertigated twice a day to keep the potting mix constantly moist. Fertigation water contained a mix of Peter’s 20-20-20 (ICL Specialty Fertilizers, Summerville, SC) and calcium nitrate (Yara) at 1.5 E.C., supplemented monthly with 50 g of the slow-release fertilizer, Osmocote plus (ICL Specialty Fertilizers, Summerville, SC). Soil electrical conductivity (E.C.) was monitored, and additional fertilizer was aqueously applied when E.C. fell below 1.2 dS m^-1^.

### 4.4. Pollination

Around thirty-six months post-transformation and 18 months after soil transplantation, the plants began to flower. The fruits in this study originated from flowers that were hand-pollinated within 24 hours of opening. To pollinate, stamens were removed using fine point tweezers, and the anthers were rubbed directly onto the stigma of the ovule donor. Each flower was pollinated with at least 2 anthers. Pollinated flowers had their stamens intact. Crossings were recorded by pressing tagged insect pins into the cacao trunk or branch near the base of the pollinated flower’s peduncle.

### 4.5. YFP Visualization and Seed Measurements

Plant tissues were visually screened for YFP presence using a Leica M165 FC stereomicroscope equipped with a 514 nm excitation and 527 nm emission filter (JH Technologies, Fremont, CA). Images were captured at 7.3-11.0x magnification using an attached Leica DFC7000 T camera and processed with Leica Application Suite X software.

To determine the inheritance pattern of transgenes, YFP expression was visualized in the seeds, radical, or first true leaves depending on the parents of the cross. To ensure a quick and high germination rate, the fruit pulp and seed coat were manually removed, and the seeds were placed into parafilm-sealed Petri dishes half-submerged in water. After two days, the seeds were washed and planted into 6.5 cm pots filled with wetted Sun Gro™ potting mix. After fourteen days, the first leaves were fully expanded, and the tip of a leaf from each plant was manually removed for YFP expression screening.

To assess whether YFP inheritance affected seed size in the progeny of Matina 1-6 EVT1, YFP expression was viewed in each seed’s emerging radicle 2 days post-germination. Seed size was measured using ImageJ, with length and width measured at the longest and widest points.

### 4.6. PCR Analysis

Genomic DNA was extracted from leaf tissue of both non-transgenic and transgenic INIAPG-038 and Matina 1-6 plants using a CTAB DNA extraction protocol as described by Murray and Thompson [40]. Progeny from the cross of INIAPG-038 EVT1 with Matina 1-6 EVT1 were tested for the transgenes CaMV35Sp*-npt*II*:eyfp* and *cas9*. The presence of CaMV35Sp-nptII was identified using the primer set 35Sp 5F (5’-CAAGTG-GATTGATGTGACATCTC-3’) and EYFP 1R (5’-TCGTCCTTGAAGAAGATGGTGC-3’), while *cas9* presence was tested with the primer set mCas9 4F (5’-CAGCGACGTGGACAAGCTGTTCAT-3’) and mCas9 5R (5’-AG-GCGTTGAACCGATCTTCCACG-3’) (Table S1). Progeny resulting from the cross of INIAPG-038 EVT2 or EVT3 with Matina 1-6 EVT1 were tested for the transgenes NOSp*-eyfp* and *cas9*. To distinguish the presence of NOSp*-eyfp* in INIAPG-038 EVT2 or EVT3 transformed with pDDNPTYFP-2 from Matina 1-6 EVT1 transformed with pMGCC3YFP, two different forward primers, oNOSp 2F and oNOS 3Fwere intentionally designed to amplify vector-specific junction regions between the sequence immediately upstream of the NOS promoter and the *eyfp* gene in each construct. the primer set oNOSp 2F (5’-TTTACGTTTGGAACTGACAGA-3’) and EYFP 1R was used for INIAPG-038 EVT2 or EVT3, while the primer set oNOSp 3F (5’-TCTAGAGGATCCCCGGGTACGA-3’) and EYFP 1R was used to identify the presence of NOSp*-eyfp* in Matina 1-6 EVT1 (Tables S1 and S2, Figures. 1 and 2). The same primer set was used for *cas9* as described above. PCR reactions were comprised of 25 μL of DreamTaq PCR Master Mix (2x), 1 μL of the 10 μM forward primer, 1 μL of the 10 μM reverse primer, 21 μL of molecular water, and 2 μl of genomic DNA at 80 ng/μL for a total reaction volume of 50 μL.

The PCR conditions for 35Sp 5F/EYFP 1R and NOSp 2F/EYFP 1R consisted of an initial denaturation at 95°C for 2 min, followed by 10 touchdown cycles of 95°C for 30 sec, annealing for 30 sec at 58°C (−0.5°C/cycle), extension at 72°C for 1.5 min, followed by 20 standard cycles of 95°C for 30 sec, annealing for 30 sec at 50°C, extension at 72°C for 40 sec, final extension at 72°C for 5 min, and 10°C for infinity. The PCR conditions for mCas9 4F/mCas9 5R followed the same protocol except for an initial touchdown annealing temperature of 65°C and a standard annealing temperature of 59°C. For each PCR reaction, 10 μL was loaded onto a 1% agarose gel for electrophoresis.

### 4.7. Copy number analysis by digital droplet PCR (ddPCR)

Transgene copy number analysis of T_0_ plants was performed using the Bio-Rad QX200 ddPCR System as described previously with minor modifications [41]. Primers used for detection of target and reference genes are described in Table 1. Copies per genome of YFP and NPTII were determined by comparing target counts to those of the reference gene TcEF1a, which we confirmed to be present as a single copy (2n) in the T. cacao genome by BLAST analysis. To prepare ddPCR reaction mixtures, purified gDNA was digested with KpnI, and 10 ng of digested template was added to each PCR reaction with 2x QX200™ ddPCR™ EvaGreen Supermix and 1 nM forward and reverse primers. Droplets were generated from 20 uL reaction mixture and 70 uL Bio-Rad Droplet Generation Oil using a following the manufacturer’s instructions. Amplification was performed with the following conditions: 10 min initial denaturation at 95 °C, 40 cycles of 1 min denaturation at 95°C and 1 min extension at 60°C with a 2°C/s ramp rate, 4°C for 5 min, 90 °C for 10 min. Droplets were analyzed using a QX200 droplet reader and the Bio-Rad QuantaSoft software using the default settings for threshold determination and quantification of positive and negative droplets. Two separate leaves were analyzed for each transgenic event, and ddPCR measurements were performed in triplicate. To compute the copy number estimations, the absolute quantification of the target was divided by that of the reference gene and multiplied by 2 to account for the two copies of EF1a in the diploid *T. cacao* genome.

### 4.8. Statistical Analysis

A **χ**^**2**^ test was used to analyze the deviation of observed segregation ratios in progeny from the expected Mendelian segregation ratios. Seed size interaction with transgene inheritance was tested using ANOVA and a Tukey-Kramer test. An α level of 0.05 was to determine significance in all statistical analyses.

## Supplementary Materials

The following supporting information can be downloaded at: www.mdpi.com/xxx/s1, Table S1: Primer sets used for selecting transgenic plants and detecting YFP expression in four different transgenic cacao events and their progeny. Table S2: Analysis of YFP expression and transgene inheritance in T_1_ progeny plants resulting from self-pollinated Matina 1-6 EVT1 and crosses between INIAPG-038 EVT1 or EVT2 and Matina 1-6 EVT1. Table S3. Primer sets used for copy number analysis by digital droplet PCR (ddPCR). Figure S1: YFP expression in T_0_ *Theobroma cacao* L. plants. Figure S2. Western blot analysis of YFP protein levels in leaves of transgenic INIAPG-038 and Matina 1-6 cacao plants.

## Author Contributions

Conceptualization, M.-J.C.; methodology, G.A., J.J. and M.-J.C.; validation and investigation, G.A., J.J., A.S., E.Z., T.T., M.G., G.V. and Y.O.; formal analysis, G.A. and M-J.C.; resources, J.-P.M. and C.J.; data curation, G.A.; writing—original draft preparation, G.A. and M.-J.C.; writing—review and editing, G.A., J.J., A.S., E.Z., T.T., M.G., J.-P.M., C.J., B.S., G.V., Y.O. and M-J.C; visualization, G.A.; supervision, M.-J.C; project administration and funding acquisition, B.S. and M-J.C. All authors have read and agreed to the published version of the manuscript.

## Data Availability Statement

Data is contained within the article.

## Acknowledgments

We thank Alexandra Tempeleu (Mars Cacao Laboratory, Miami, FL) for kindly providing *Theobroma cacao* flowers, as well as Naio Koehler and Christina Wistrom (University of California, Berkeley, CA) for their dedicated care of the plants in the growth chambers and greenhouse.

## Disclaimer/Publisher’s Note

The statements, opinions and data contained in all publications are solely those of the individual author(s) and contributor(s) and not of MDPI and/or the editor(s). MDPI and/or the editor(s) disclaim responsibility for any injury to people or property resulting from any ideas, methods, instructions or products referred to in the content.

## Supplementary Information

**Table S1.**
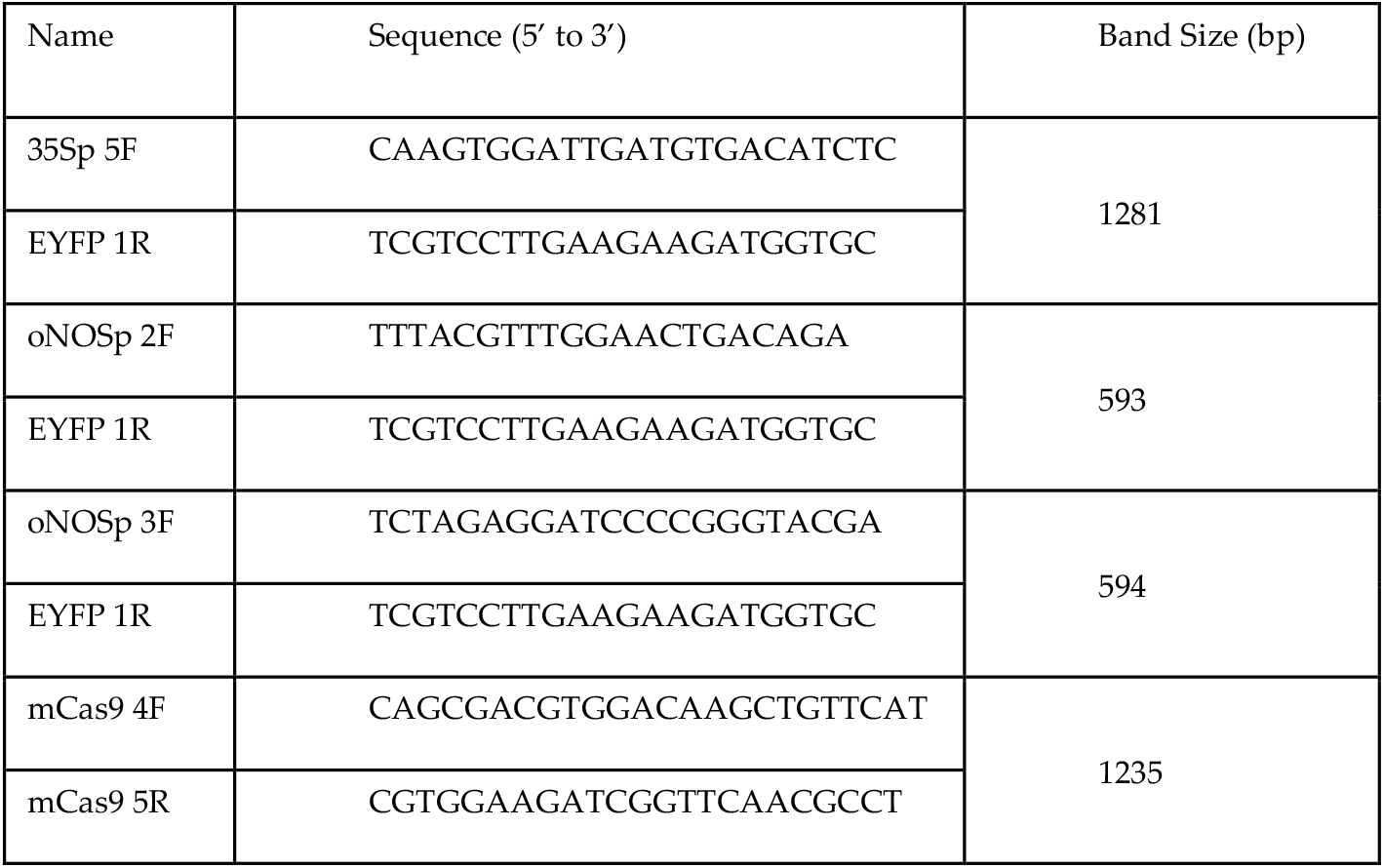
Primer sets used for selecting transgenic plants and detecting YFP expression in four different transgenic cacao events and their progeny.

**Table S2.**
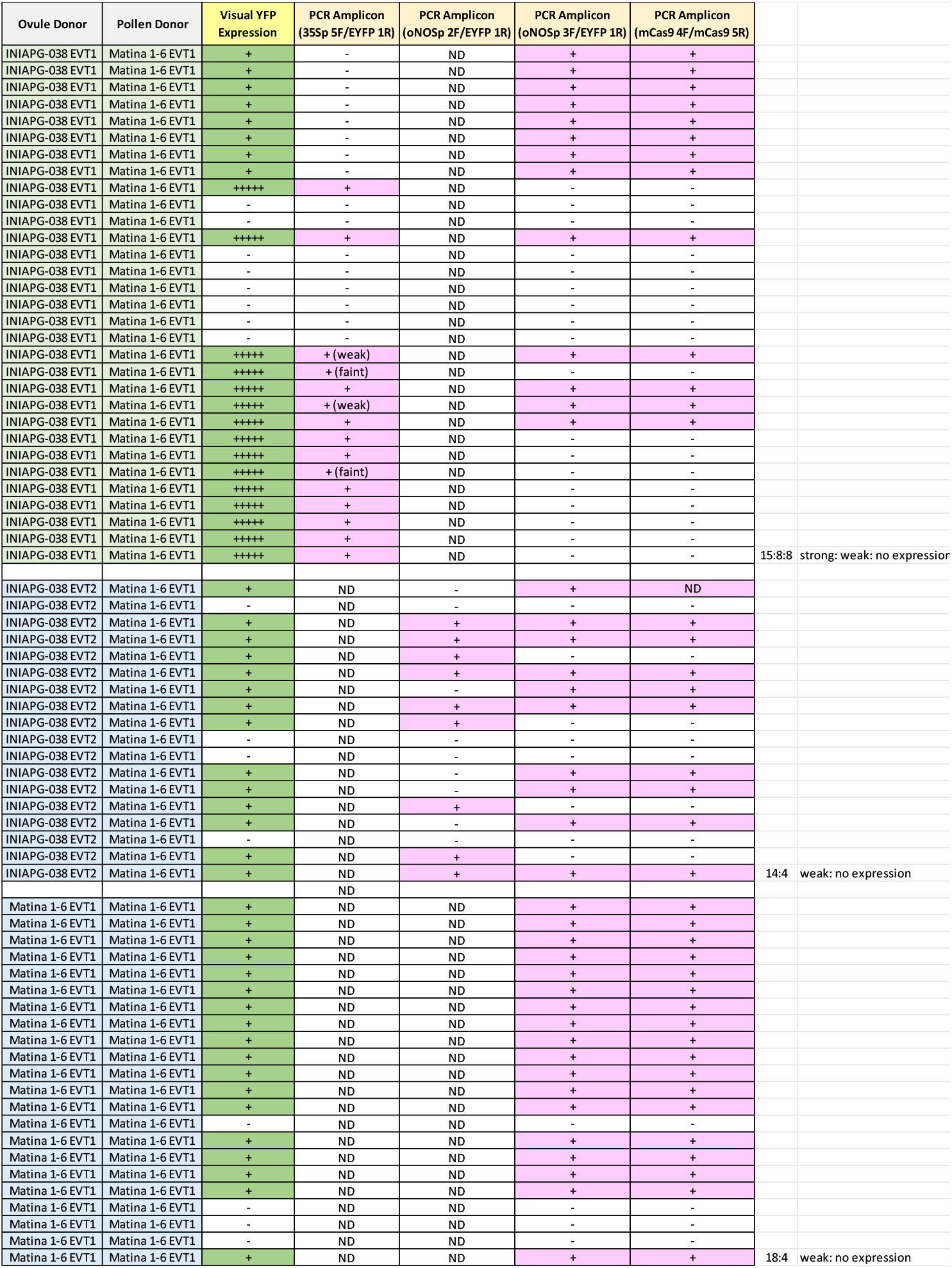
Analysis of YFP expression and transgene inheritance in T_1_ progeny plants resulting from self-pollinated Matina 1-6 EVT1 and crosses between INIAPG-038 EVT1 or EVT2 and Matina 1-6 EVT1. Reciprocal crosses between INIAPG-038 EVT1 and Matina 1-6 EVT1 yielded a 2:1:1 segregation (strong: weak: no expression), consistent with the expected 3:1 positive-to-negative ratio. The strong CaMV 35S–driven YFP in INIAPG-038 EVT1 masked the weaker NOSp::TMV Ω–driven signal, making progeny inheriting both transgenes indistinguishable from those carrying only the CaMV 35S cassette. No moderately strong YFP expression was observed in T1 progeny plants.

**Table S3.**
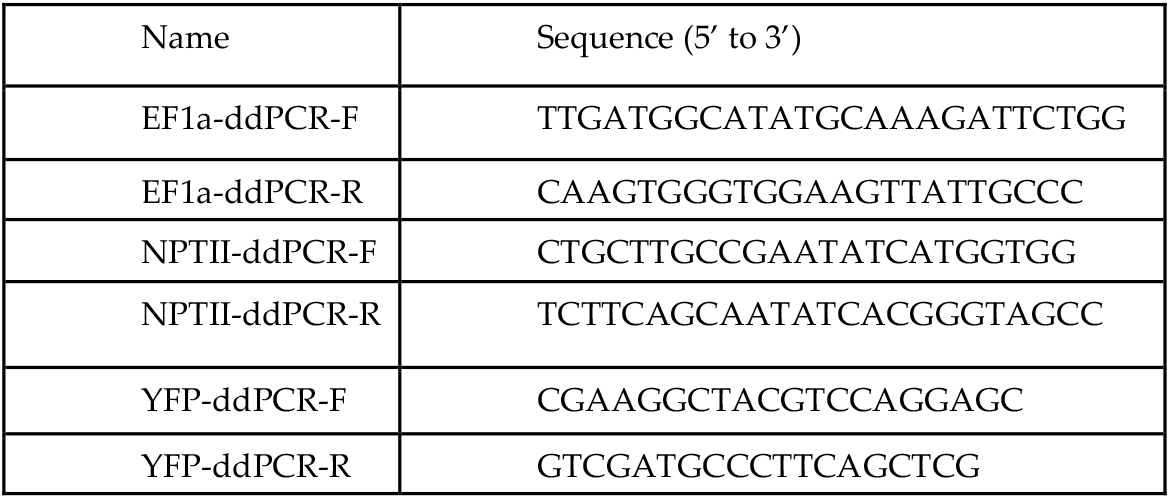
Primer sets used for copy number analysis by digital droplet PCR (ddPCR).

**Figure S1.**
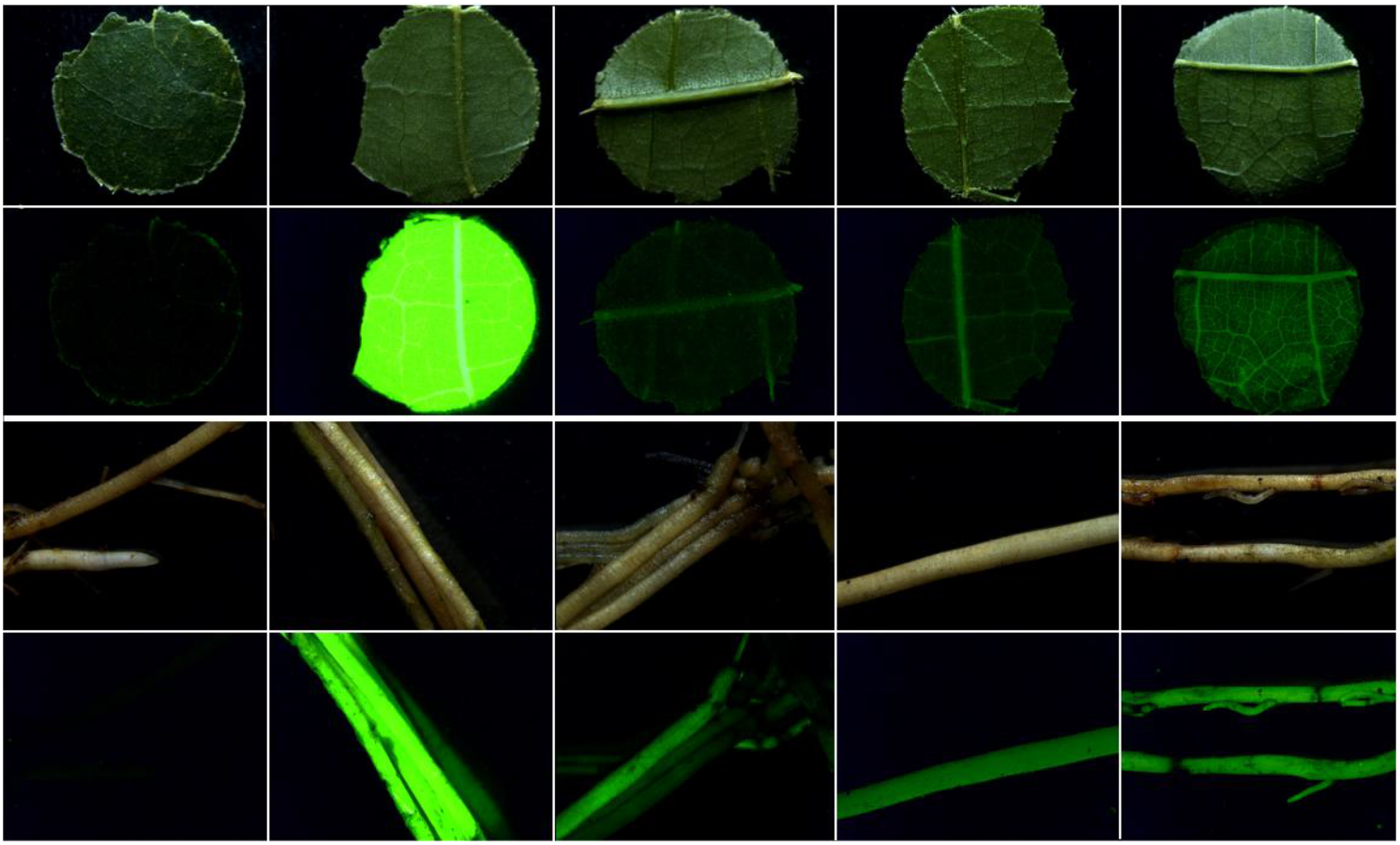
YFP expression in T_0_ *Theobroma cacao* L. plants. (A) Top row and third row: YFP filter; second row and bottom row: brightfield. From left to right: leaf and root samples from non-transgenic INIAPG-038, INIAPG-038 EVT 1, INIAPG-038 EVT 2, INIAPG-038 EVT 3, and Matina 1-6 EVT1.

**Figure S2.**
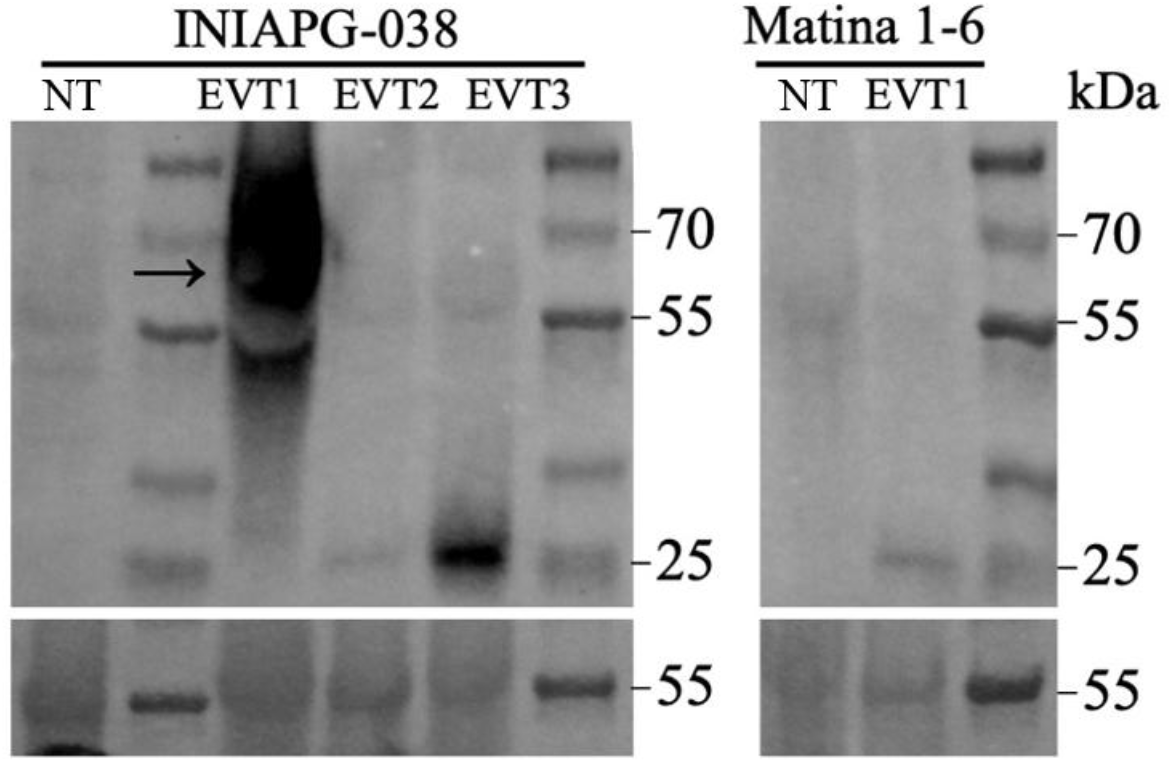
Western blot analysis of YFP protein levels in leaves of transgenic INIAPG-038 and Matina 1-6 cacao plants. The upper panel shows immunodetection of YFP in young leaf samples from six cacao plants using an anti-YFP antibody. The lower panel shows the corresponding Rubisco large subunit signal as a loading control. Samples include one non-transgenic INIAPG-038 plant (NT), three independent transgenic INIAPG-038 events (EVT1, EVT2 and EVT3), one non-transgenic Matina 1-6 plant (NT), and one transgenic Matina 1-6 plant (EVT1). YFP expression was detected only in the transgenic lines.

### Western Blot Analysis of YFP Expression Levels

Immunoblot assays were performed as previously described (Qi et al., 2018), with minor modifications. Briefly, total protein was extracted from cacao leaves using 2× Laemmli buffer containing 5% β-mercaptoethanol. Proteins were separated on a 4–15% precast polyacrylamide gel (Bio-Rad, Hercules, CA), transferred to nitrocellulose membranes, and then probed with a mouse anti-YFP antibody (1:1,000; Biosensis Pty Ltd., Thebarton, SA, Australia). Antigens were detected using the SuperSignal™ West Pico PLUS Chemiluminescent Substrate (Thermo Fisher Scientific, Waltham, MA) and imaged with a ChemiDoc XRS+ system (Bio-Rad, Hercules, CA). Ponceau S staining was used as a loading control.

## References

1. Abt, E.; Robin, L.P. Perspective on cadmium and lead in cocoa and chocolate. J. Agric. Food Chem. 2020, 68, 13008–13015.

2. Motamayor, J.C., Lachenaud, P.; da Silva e Mota JW, Loor, R.; Kuhn, D.N.; Brown, J.S.; Schnell, R.J. (2008). Geographic and genetic population differentiation of the Amazonian chocolate tree (Theobroma cacao L.). PLOS ONE, 2008, 3, e3311.

3. Bustamante, D.E.; Motilal, L.A.; Calderon, M.S.; Mahabir, A.; Oliva, M. Genetic diversity and population structure of fine aroma cacao (Theobroma cacao L.) from north Peru revealed by single nucleotide polymorphism (SNP) markers. Front. Ecol. Evol. 2022, 10, 895056.

4. Cornejo, O.E.; Yee, M.C.; Dominguez, V.; Andrews, M.; Sockell, A.; Strandberg, E.; Motamayor, J.C. Population genomic analyses of the chocolate tree, Theobroma cacao L., provide insights into its domestication process. Commun. Biol. 2018, 1, 167.

5. Marelli, J.P.; Guest, D.I.; Bailey, B.A.; Evans, H.C.; Brown, J.K.; Junaid, M.; Barreto, R.W.; Lisboa, D.O.; Puig, A.S. Chocolate under threat from old and new cacao diseases. Phytopathology 2019, 109, 1331–1343.

6. Mulhollem, J. Discovery of flowering gene in cacao may lead to accelerated breeding strategies. Penn State 2021. https://www.psu.edu/news/research/story/discovery-flowering-gene-cacao-may-lead-accelerated-breeding-strategies/.

7. Gotsch, N. Cocoa biotechnology: status, constraints and future prospects. Biotechnol. Adv. 1997, 15, 333–352.

8. Wickramasuriya, A.M.; Dunwell, J.M. Cacao biotechnology: current status and future prospects. Plant Biotechnol. J. 2018, 16, 4– 17.

9. Bailey-Serres, J.; Parker, J.E.; Ainsworth, E.A.; Oldroyd, G.E.; Schroeder, J.I. Genetic strategies for improving crop yields. Nature 2019, 575, 109–118.

10. Zhang, Y.; Malzahn, A.A.; Sretenovic, S.; Qi, Y. The emerging and uncultivated potential of CRISPR technology in plant science. Nat. Plants 2019, 5, 778–794.

11. Maximova, S.; Miller, C.; Antunez de Mayolo, G.; Pishak, S.; Young, A.; Guiltinan, M. J. Stable transformation of Theobroma cacao L. and influence of matrix attachment regions on GFP expression. Plant Cell Rep. 2003, 21, 872–883.

12. Jones, J.; Zhang, E.; Tucker, D.; Rietz, D.; Dahlbeck, D.; Gomez, M.; Garcia, C.; Marelli, J.-P.; Livingston III, D.; Schnell, R.; et al. Screening of cultivars for tissue culture response and establishment of genetic transformation in a high-yielding and disease-resistant cultivar of Theobroma cacao. In Vitro Cell. Dev. Biol.-Plant 2022, 58, 133–145.

13. McElroy, M.S.; Navarro, A.J.; Mustiga, G.; Stack, C.; Gezan, S.; Peña, G.; Sarabia, W.; Saquicela, D.; Sotomayor, I.; Douglas, G.M.; et al. Prediction of cacao (Theobroma cacao) resistance to Moniliophthora spp. diseases via genome-wide association analysis and genomic selection. Front. Plant Sci. 2018, 9, 343.

14. Meinhardt, L.W.; Rincones, J.; Bailey, B.A.; Aime, M.C.; Griffith, G.W.; Zhang, D.; Pereira, G.A. Moniliophthora perniciosa, the causal agent of witches’ broom disease of cacao: What’s new from this old foe? Mol. Plant Pathol. 2008, 9, 577–588.

15. Phillips-Mora, W.; Baqueros, F.; Melnick, R.L.; Bailey, B.A. First report of frosty pod rot caused by Moniliophthora roreri on cacao in Bolivia. New Dis. Rep. 2015, 31, 29.

16. Motamayor, J.C.; Mockaitis, K.; Schmutz, J.; Haiminen, N.; Livingstone III, D.; Cornejo, O.; Findley, S.D.; Zheng, P.; Utro, F.; Royaert, S.; et al. The genome sequence of the most widely cultivated cacao type and its use to identify candidate genes regulating pod color. Genome Biol. 2013, 14, r53.

17. Ofori, A.; Padi, F.K.; Ameyaw, G.A.; Dadzie, A.M.; Opoku-Agyeman, M.; Domfeh, O.; Ansah, F.O. Field evaluation of the impact of cocoa swollen shoot virus disease infection on yield traits of different cocoa (Theobroma cacao L.) clones in Ghana. PLoS One 2022, 17, e0262461.

18. Lee, M.; Phillips, R.L. The chromosomal basis of somaclonal variation. Annu. Rev. Plant Physiol. Plant Mol. Biol. 1988, 39, 413– 437.

19. Phillips, R.L.; Kaeppler, S.M.; Olhoft, P. Genetic instability of plant tissue cultures: breakdown of normal controls. Proc. Natl. Acad. Sci. 1994, 91, 5222–5226.

20. Dietz-Pfeilstetter, A. Stability of transgene expression as a challenge for genetic engineering. Plant Sci. 2010, 179, 164–167.

21. Vaucheret, H.; Béclin, C.; Elmayan, T.; Feuerbach, F.; Godon, C.; Morel, J.-B.; Mourrain, P.; Palauqui, J.-C.; Vernhettes, S. Transgene-induced gene silencing in plants. Plant J. 1998, 16, 651–659.

22. Laboulaye, M.A.; Duan, X.; Qiao, M.; Whitney, I.E.; Sanes, J.R. Mapping transgene insertion sites reveals complex interactions between mouse transgenes and neighboring endogenous genes. Front. Mol. Neurosci. 2018, 11, 385.

23. Baulcombe, D. RNA silencing in plants. Nature 2004, 431, 356–363.

24. Meyer, P. Stabilities and instabilities in transgene expression. In Transgenic Plant Research; Routledge: 1998; pp. 263–275.

25. Sena Gomes, A.R.; Andrade Sodré, G.; Guiltinan, M.; Lockwood, R.; Maximova, S. Supplying new cocoa planting material to farmers: A Review of Propagation Methodologies; Laliberte, B.; End, M., Eds.; Bioversity International. Rome, Italy, 2015.

26. Privalle, L.; Back, P.; Bhargava, A.; Bishop, Z.; Cisneros, K.; Coats, I.; Criel, I.; Dhondt, L.; Draughn, T.; Fowler, B.; et al. Genetic stability, inheritance patterns and expression stability in biotech crops. OBM Genet. 2020, 4, 1.

27. Sanders, P.R.; Winter, J.A.; Barnason, A.R.; Rogers, S.G.; Fraley, R.T. Comparison of cauliflower mosaic virus 35S and nopaline synthase promoters in transgenic plants. Nucleic Acids Res. 1987, 15, 1543–1558.

28. Horstmann, V.; Huether, C.M.; Jost, W.; Reski, R.; Decker, E.L. Quantitative promoter analysis in Physcomitrella patens: a set of plant vectors activating gene expression within three orders of magnitude. BMC Biotechnol. 2004, 4, 1–13.

29. Mannerlöf, M.; Tenning, P. Variability of gene expression in transgenic tobacco. Euphytica, 1997, 98, 133–139.

30. Glendinning, D.R. Natural pollination of cocoa. New Phytol. 1972, 71, 719–729.

31. Lanaud, C.; Fouet, O.; Legavre, T.; Lopes, U.; Sounigo, O.; Eyango, M.C.; Mermaz, B.; Da Silva, M.R.; Loor Solozano, R.G.; Argout, X.; et al. Deciphering the Theobroma cacao self-incompatibility system: from genomics to diagnostic markers for self-compatibility. J. Exp. Bot. 2017, 68, 4775–4790.

32. Choi, H.-W.; Lemaux, P.G.; Cho, M.-J. High-frequency of cytogenetic aberration in transgenic oat (Avena sativa L.) plants. Plant Sci. 2001, 160, 763–772.

33. Cho, M.-J.; Choi, H.-W.; Bregitzer, P.; Zhang, S.; Lemaux, P.G. Transgenic barley (Hordeum vulgare L.) and chromosomal variation. In Testing for Genetic Manipulation in Plants. Molecular Methods of Plant Analysis; Jackson, J.F., Linskens, H.F., Inman, R.B., Eds.; Springer Verlag. 2002; volume 22, pp. 169–188.

34. Farinati, S.; Draga, S.; Betto, A.; Palumbo, F.; Vannozzi, A.; Lucchin, M.; Barcaccia, G. Current insights and advances into plant male sterility: new precision breeding technology based on genome editing applications. Front. Plant Sci. 2023, 14, 1223861.

35. Kozaki, A.; Aoyanagi, T. Molecular aspects of seed development controlled by gibberellins and abscisic acids. Int. J. Mol. Sci. 2022, 23, 1876.

36. Matilla, A.J. Auxin: hormonal signal required for seed development and dormancy. Plants. 2020, 9, 705.

37. Lebedev, V. Stability of transgene inheritance in progeny of field-grown pear trees over a 7-year period. Plants 2022, 11, 151.

38. Batista, R.A.; Köhler, C. Genomic imprinting in plants—revisiting existing models. Genes Dev. 2020, 34, 24–36.

39. Gomez, M.A.; Berkoff, K.C.; Gill, B.K.; Iavarone, A.T.; Lieberman, S.E.; Ma, J.M.; Schultink, A.; Karavolias, N.G.; Wyman, S.K.; Chauhan, R.D.; et al. CRISPR-Cas9-mediated knockout of CYP79D1 and CYP79D2 in cassava attenuates toxic cyanogen production. Front. Plant Sci. 2023, 13, 1079254.

40. Murray, M.G.; Thompson, W.F. Isolation of plant DNA from fresh tissue. Nucleic Acids Res.1980, 8, 4321–4325.

41. Collier, R; Dasgupta, K.; Xing, YP, Hernandez BT, Shao M, Rohozinski D, Kovak E, Lin J, de Oliveira ML, Stover E, McCue KF. Accurate measurement of transgene copy number in crop plants using droplet digital PCR. Plant J. 2017, 90, 1014–25.

## Reference

Qi, T., Seong, K., Thomazella, D.P.T., Kim, J.R., Pham, J., Seo, E., Cho, M.-J., Schultink, A., Staskawicz, B.J. NRG1 functions downstream of EDS1 to regulate TIR-NLR-mediated plant immunity in Nicotiana benthamiana, Proc. Natl. Acad. Sci. 2018, 115, E10979-E10987.

